# Predicting antibiotic-associated virulence of *Pseudomonas aeruginosa* using an *ex-vivo* lung biofilm model

**DOI:** 10.1101/2020.02.24.963173

**Authors:** Marwa M. Hassan, Niamh E. Harrington, Esther Sweeney, Freya Harrison

## Abstract

**Background:** Bacterial biofilms are known to have high antibiotic tolerance which directly affects clearance of bacterial infections in people with cystic fibrosis (CF). Current antibiotic susceptibility testing methods are either based on planktonic cells or do not reflect the complexity of biofilms *in vivo*. Consequently, inaccurate diagnostics affect treatment choice, preventing bacterial clearance and potentially selecting for antibiotic resistance. This leads to prolonged, ineffective treatment.

**Methods:** In this study, we use an *ex-vivo* lung biofilm model to study antibiotic tolerance and virulence of *Pseudomonas aeruginosa*. Sections of pig bronchiole were dissected, prepared and infected with clinical isolates of *P. aeruginosa* and incubated in artificial sputum media to form biofilms, as previously described. Then, lung-associated biofilms were challenged with antibiotics, at therapeutically relevant concentrations, before their bacterial load and virulence were quantified and detected, respectively.

**Results:** The results demonstrated minimal effect on the bacterial load with therapeutically relevant concentrations of ciprofloxacin and meropenem, with the later causing an increased production of proteases and pyocyanin. A combination of meropenem and tobramycin did not show any additional decrease in bacterial load but demonstrated a slight decrease in total proteases and pyocyanin production.

**Conclusions:** We demonstrate a realistic model for understanding antibiotic resistance and tolerance in biofilms clinically and for molecules screening in anti-biofilm drug development. *P. aeruginosa* showed high levels of antibiotic tolerance, with minimal effect on bacterial load and increased proteases production, which could negatively affect lung function. This may potentially contribute to exacerbations and eventual lung failure.

## Introduction

Cystic fibrosis (CF) is a genetic disease in which people have decreased mucociliary clearance in the respiratory tract, due to mutations in the cystic fibrosis transmembrane conductance regulator (*CFTR*) gene, which encodes a chloride channel [1, 2]. This impairment leads to a reduction in mucus clearance and increased viscosity, resulting in accumulation of inhaled microbial cells, increased bacterial adherence and inflammation and the formation of bacterial biofilm [1-3]. Biofilm infections are more difficult to eradicate due the difference in their nature compared to non-biofilm infections; thus, they are lifelong infections in CF. These biofilm infections are characterised by acquiring distinctive resistance mechanisms compared with non-biofilm infections. There are three main mechanisms. First, there may be low antibiotic penetration into the biofilm due to the production of extracellular matrix. Second, the different bacterial metabolic states in the biofilm lead to increased phenotypic heterogeneity, affecting the success of treatment. Third, adaptive mechanisms controlling differential gene expression of multiple virulence factors, such as efflux pumps and antibiotic-degrading enzymes, lead to antibiotic tolerance [4, 5]. The latter is highly dependent and varies based on the environment surrounding the biofilm [5].

In CF and other biofilm-based infections, antibiotic prescription is mainly based on standard minimum inhibitory concentration (MIC) methods, despite these being based on planktonic cells [6]. Planktonic-based diagnostics are suitable for detecting intrinsic and acquired stable resistance mechanisms; however, biofilm-based diagnostics will additionally detect environmentally-induced and biofilm-associated resistance mechanisms. There is a drastic increase in antibiotic tolerance using biofilm-based diagnostics, such as the Calgary device, in comparison to MIC methods, demonstrating the limitation of using planktonic-based models for biofilm infections [7]. Some of these *in vitro* biofilm models are robust for antibiotic susceptibility screening; they fail to recapitulate the complexity of biofilm infections, environment and host-dependent interactions [6, 8], all of which affect the antibiotic susceptibility profile [8]. Also, these biofilm models have not been developed to demonstrate antibiotic susceptibility profile in multi-species infections. Thus, if diagnostic tests fail to accurately detect *in vivo* antibiotic resistance, this will result in in recurrent and complicated infections [8], and may lead to a vicious cycle of increased resistance.

In this study, we employed a previously developed *ex-vivo* pig lung biofilm model (EVPL) [9, 10] for antibiotic susceptibility testing of CF *P. aeruginosa* isolates and compared it with standard MIC and the Calgary device assays. The effect of exposure to antibiotics (ciprofloxacin, meropenem, tobramycin and combination therapy) on the virulence of *P. aeruginosa* was also assessed. The results demonstrated an increased antibiotic tolerance in EVPL model at >25-fold the reported sputum concentrations when tested in Mueller-Hinton broth, which even further increased when tested in artificial sputum media. Exposure to antibiotics showed an increased production of total proteases, which may have a role in lung damage. Normalised proteases/cfu and pyocyanin/cfu demonstrated an increased production of these virulence factors per cell. Current clinical prognosis in CF is alarming and creates an urgent need to develop effective anti-biofilm agents. Thus, we propose a unique approach to predict the true clinical effect of antibiotic treatments on bacterial clearance and associated virulence factors in CF using the EVPL model.

## Methods

### 1) Bacterial isolates

Clinical *Pseudomonas aeruginosa* isolates used in this study were isolated from a chronically-colonised CF adult, as described in Darch *et al.* [11]. Six bacterial isolates were chosen to represent a range of different growth rates and virulence profiles as reported by Darch *et al.* [11]. PA14 was used as a control laboratory strain for comparison.

### 2) Antibiotic susceptibility testing

Minimum inhibitory concentrations were performed according to the EUCAST guidelines [12]. Briefly, bacterial isolates were cultured in LB agar overnight at 37°C, resuspended in Mueller-Hinton broth (MHB) to OD_600_ of 0.5 and diluted 1000 times. Meropenem (Sigma Aldrich) and ciprofloxacin (Thermo Fisher) were two-fold serially diluted in MHB, to a final volume of 50 µL in a 96-well plate (Corning). 50 µL of the diluted bacterial suspension (5×10^5^ cfu/mL final concentration) was added to all wells and incubated for 18 hours at 37°C.

Minimum biofilm eradication concentrations (MBEC) were performed according to Moskowitz *et al.* [13]. Briefly, 100 µL of bacterial suspensions at 0.5 McFarland were aliquoted in U shaped 96-well plates (Corning), covered with peg lids (Thermo Electron) and incubated for 20 hours at 37°C to form biofilms on the pegs. Peg lids were then washed three times in sterile phosphate-buffered saline (PBS), transferred to 96-well antibiotic challenge plates containing 100 µL of 2-fold serially diluted meropenem or ciprofloxacin and incubated for 18 hours at 37°C. Peg lids were washed three times in sterile PBS, transferred to 96-well recovery plates containing 100 µL MHB and sonicated for 5 minutes. Peg lids were replaced with standard plate lids and incubated for 6 hours at 37°C before checked for turbidity.

### 3) Antibiotic tolerance in EVPL

Dissection and infection of pig lungs were performed as described in Harrison *et al.* [9]. Briefly, pig bronchioles were dissected, UV sterilised and transferred to 24-well plates with 400 µL of 0.8% agarose/ASM (Artificial Sputum Medium [14]) as a pad. Bronchiole tissues were infected with the bacterial isolates using a syringe and 500 µL of ASM were added to each well. Uninfected bronchiole tissues were used as negative controls. Plates were then covered with UV sterilised breathable membranes (Sigma Aldrich) and incubated at 37°C for 7 days. Tissues were then washed in 500 µL of PBS, transferred to sterile bead tubes containing 1 gm of metal beads (2.4 mm, Fisher Scientific) and 1 mL of PBS, and homogenised using a FastPrep-24™ 5G homogeniser (MP Biomedicals) for 40 sec at 4.0 m/sec. Biofilm homogenate was transferred to 96-well plates, serially diluted and plated on LB agar plates for calculating the bacterial load.

Replicate infected tissues were treated with antibiotics by transferring the washed infected tissues to 48-well plates containing 300 µL of either MHB or ASM containing ciprofloxacin (Thermo Fisher), meropenem (Sigma), tobramycin (Thermo Fisher) or combination therapy at the specified concentrations and incubated at 37°C for 24 hours. Antibiotic-treated tissues were then washed, homogenised and plated as previously described.

### 4) Virulence assays

Total proteases were quantified according to Harrison *et al.* [10]. Briefly, 100 µL of tissue homogenate or surrounding ASM were added to 900 µL of azocasein solution (final concentration of 5 mg/mL dissolved in 100 mM Tris-HCl, 1 mM CaCl_2_) in a 2 mL tube and incubated for 15 min at 37°C with shaking at 170 rpm. Then, 500 µL of 10% trichloroacetic acid were added as a stopping solution, and tubes were centrifuged at 13,000 rpm for 1 min at room temperature. 200 µL of the supernatant were transferred into a clean 96-well plate and the absorbance was measured at 400 nm. PBS was used as a negative control and a standard curve using proteinase K (**Figure S1**) was used to estimate the total amount of proteases.

Total pyocyanin was quantified according to Saha *et al.* [15] with minor modifications. Briefly, pyocyanin was extracted using chloroform in a ratio of 5:3. The chloroform mixture was vortexed for 2 minutes, then centrifuged at room temperature at 10,000 rpm for 5 minutes. The bottom layer was transferred to a new 2 mL tube and an equal volume of 0.2M HCl was added. Tubes were vortexed for 2 minutes, centrifuged at room temperature at 10,000 rpm for 5 minutes and 200 µL of the top phase was transferred to a black 96-well plate and the absorbance was measured at 520 nm [15]. The concentration of pyocyanin (µg/mL) was calculated by multiplying the OD_520_ by 17.072 [16].

### 5) Statistical analyses

All data were analysed by ANOVA to test for the main effect and interactions of different lung, strains and antibiotic treatments using RStudio v1.1.463 (©2009-2018 RStudio, Inc.). Unpaired t-tests were performed for pairwise statistical analysis using GraphPad Prism (v8.0.1) and the familywise error rate correction for multiple comparisons.

## Results

### 1) Antibiotic susceptibility testing

Clinical *P. aeruginosa* strains were resistant to ciprofloxacin (MIC 2 µg/mL) and sensitive to meropenem (MIC ≤0.25 µg/mL) by standard antibiotic susceptibility testing using the broth dilution method; PA14 was sensitive to both antibiotics (**Table 1**). Determination of MBECs using the Calgary device demonstrated an increase of 2–4-fold and 64–128-fold MIC for ciprofloxacin and meropenem, respectively (**Table 1**). Additionally, the MBEC recovery plates showed no visible production of pyocyanin, pyochelin or pyoverdine (**Figure S2**).

**Table 1.** Determination of MIC and MBEC of *P. aeruginosa* isolates against ciprofloxacin and meropenem.

| Strains | MIC ($\mu\text{g/mL}$)<br>(n=4) | | MBEC ($\mu\text{g/mL}$)<br>(n=4) | |
| --- | --- | --- | --- | --- |
| | Ciprofloxacin<br>$S \leq 0.5, R > 0.5$ | Meropenem<br>$S \leq 2, R > 8$ | Ciprofloxacin | Meropenem |
| SED 20 | 2 | 0.0625 | 8 | 8 |
| SED 29 | 2 | 0.25 | 4 | 16 |
| SED 34 | 2 | 0.25 | 4 | 16 |
| SED 38 | 2 | 0.0625 | 4 | 8 |
| SED 41 | 2 | 0.125 | 4 | 16 |
| SED 43 | 2 | 0.125 | 8 | 16 |
| PA14 | 0.25 | 0.25 | 1 | 0.5 |

### 2) Increased antibiotic tolerance of bacterial biofilms in the EVPL

**Figure 1** represents a schematic diagram of the work flow as previously described. **Figure S3** and **Figure S4** show pieces of tissues infected with *P. aeruginosa* strains after 7 days of biofilm formation and the mucoid phenotype of the strains in the tissues, respectively, to represent chronic CF infection [17].

The effect of antibiotics on the bacterial load was first tested by transferring bronchiole sections containing developed biofilms to MHB (standard medium for microdilution assays) containing antibiotics. This allowed to directly compare the inhibitory effect of these antibiotics in EVPL versus in standard diagnostics assays, without the effect of the medium or any physiological differences induced by CF lung mucus. Treatment with ciprofloxacin at 32-fold MIC (8–16-fold MBEC) resulted in a 1-2 log decrease in bacterial load across all tested clinical strains (**Figure 2A**). However, exposure to meropenem at 256–1024-fold MIC (4–8-fold MBEC) resulted in only about 1 log decrease of the bacterial load across all strains except SED 34, which showed less than a log decrease in the bacterial count (**Figure 2B**). ANOVA showed a significant effect of ciprofloxacin (F_1,69_ = 749.5, p < 0.001) and meropenem treatment (F_1,69_ = 115.5, p < 0.001) on bacterial load, the magnitude of which was strain dependent (strain*treatment interaction F_6,69_ = 3.1, p <0.01) for ciprofloxacin but strain independent for meropenem (strain*treatment interaction F_6,69_ = 0.71, p = 0.64) despite the differences in MICs and MBECs.

**Figure 1.**
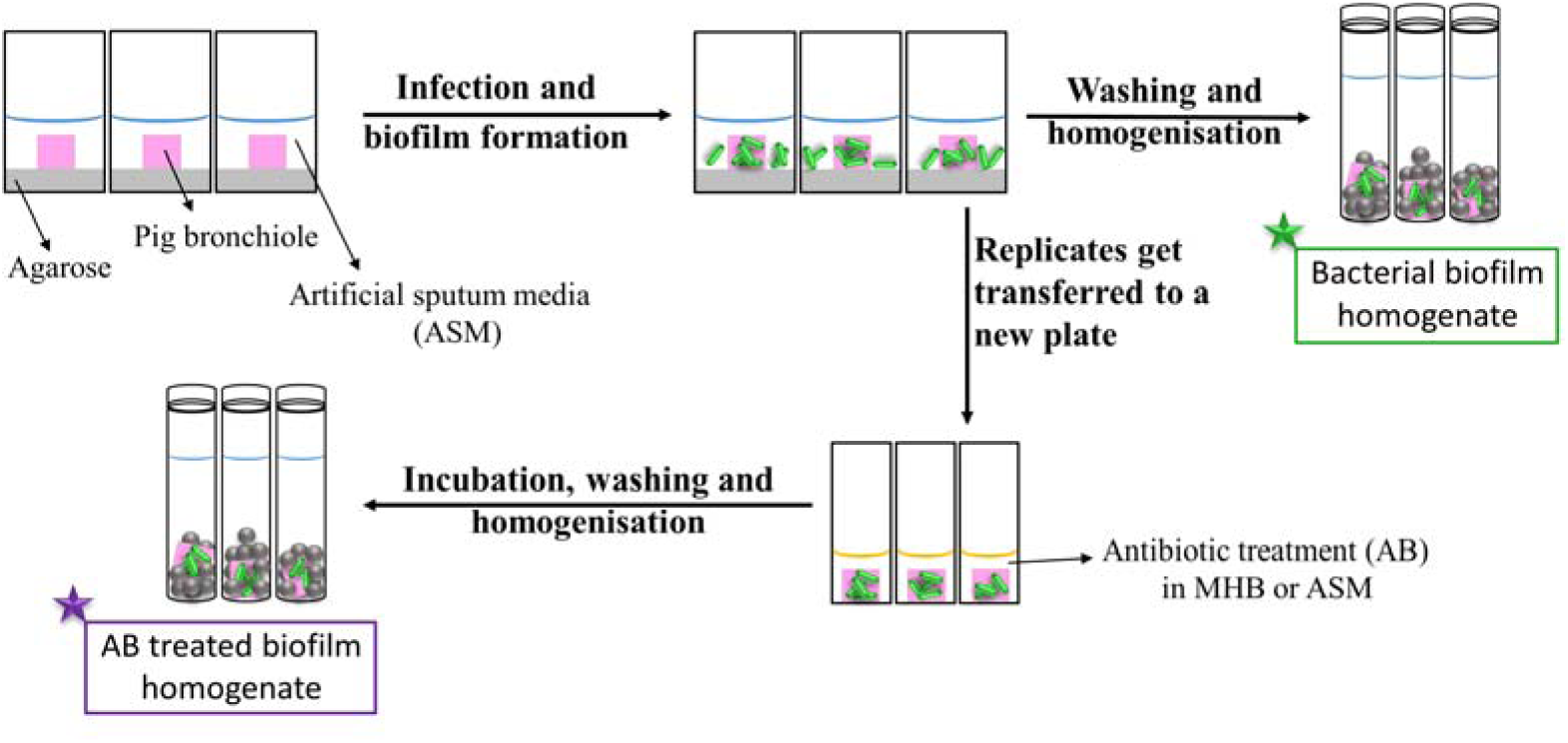
Schematic diagram of the work flow for the determination of the antibiotic susceptibility. Pig bronchioles were infected with *P. aeruginosa* clinical strains, incubated to form biofilms and homogenised for the determination of the biofilm bacterial load. Replicate infected tissues were exposed to antibiotics for 24 hours before the decrease in bacterial load was determined.

**Figure 2.**
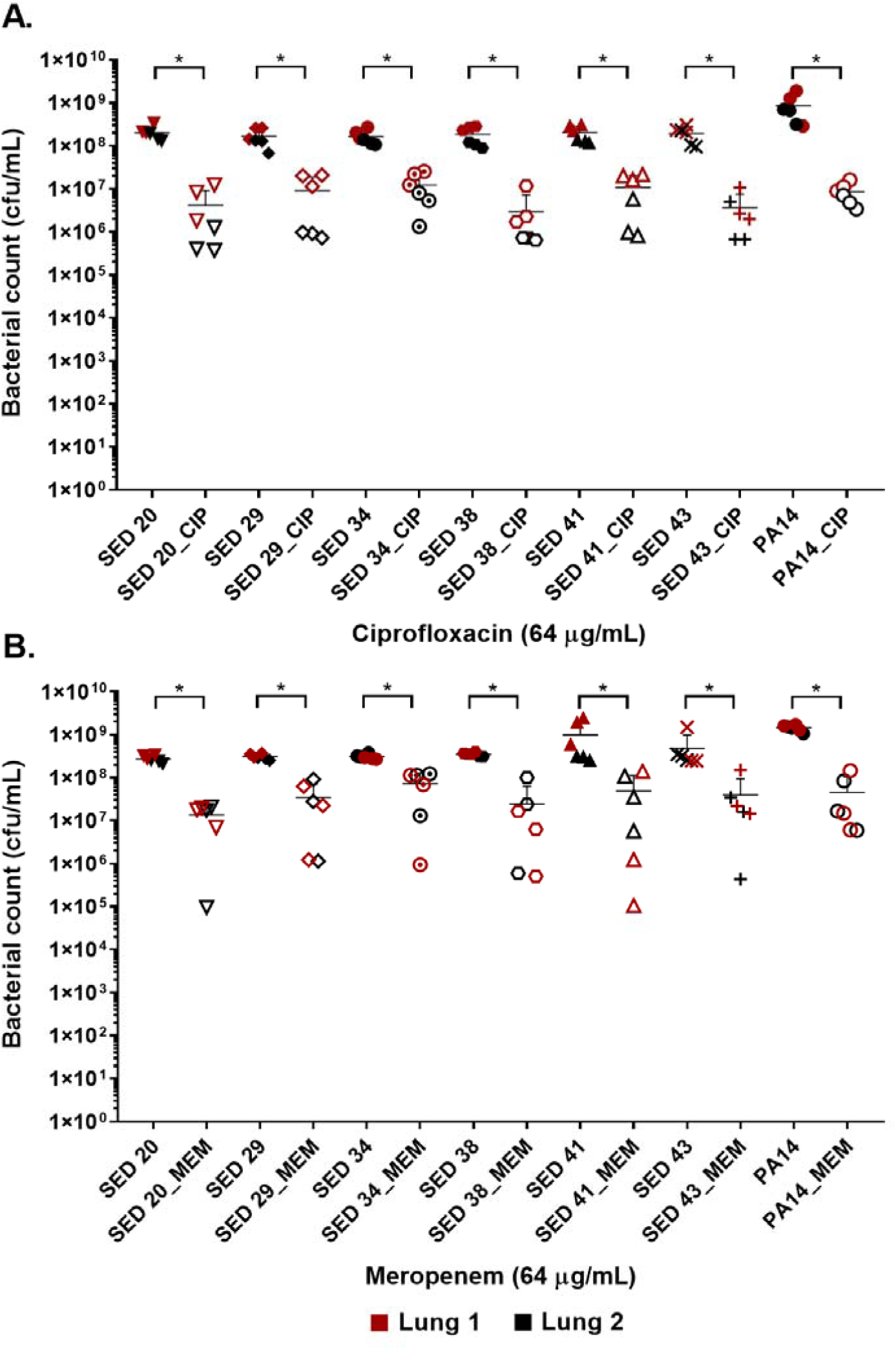
Bacterial load of *P. aeruginosa* in the EVPL biofilm model with and without exposure to A. ciprofloxacin (CIP), B. meropenem (MEM) 64 (µg/mL). Error bars are means ± SD, some error bars are too small to be visible on the graph. Unpaired t-tests were performed for the pairwise statistical analysis of treated against untreated bacterial biofilm load for each strain; significant difference (p value < 0.05) are denoted with *.

To investigate the effect of the environment on antibiotic tolerance, the bacterial load of the clinical isolate SED 43 was compared with and without ciprofloxacin or meropenem in MHB and ASM: a chemically defined medium which mimics the chemistry of chronically-infected CF sputum [14] (**Figure 3**). As shown in **Figure 3**, there was a statistically significant increase in tolerance to both ciprofloxacin and meropenem treatment in ASM (0.84 and 0.54 log decrease) compared with MHB (1.68 and 1.02 log decrease). ANOVA showed significant effects of changing of medium (F_1,20_ = 9.04, p < 0.01), antibiotic treatment (F_1,20_ = 44.66, p < 0.001) and a medium*antibiotic interaction F_1,20_ = 5.56, p < 0.05.

**Figure 3.**
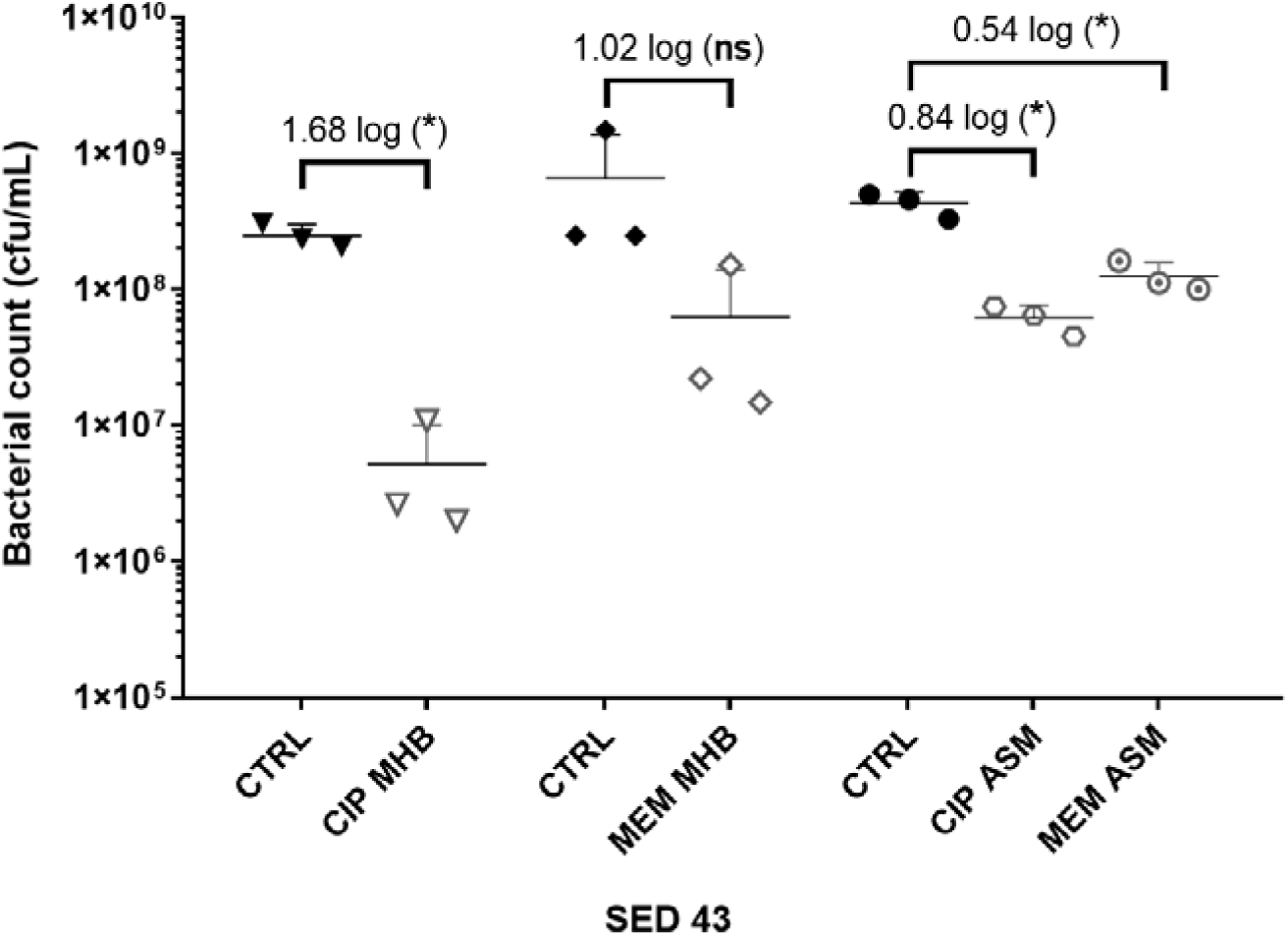
The effect of medium used (MHB or ASM) on antibiotic susceptibility of the clinical isolate SED 43 after exposure to ciprofloxacin (CIP) or meropenem (MEM) at 64 (µg/mL) in the EVPL biofilm. CTRL are controls of non-treated tissues from the same lung. Error bars are means ± SD, some error bars are too small to be visible on the graph. Unpaired t-tests were performed for the pairwise statistical analysis of treated against untreated bacterial biofilm load; significant difference (p value < 0.05) are denoted with *.

### 3) The effect of antibiotic-mediated virulence

To understand the effect of antibiotics on the virulence of *P. aeruginosa* strains, PA14 and three clinical isolates (SED 20, SED 41 and SED 43) were assessed for the production of proteases and pyocyanin in the lung tissues and surrounding ASM, separately, under exposure to antibiotics. The ASM surrounding tissue sections at the end of antibiotic exposure remained visibly clear, suggesting that bacterial cells did not detach from the biofilm or grow in the surrounding ASM at appreciable rates. Total proteases and pyocyanin were shown to be mainly released from the biofilm into the ASM, with a similar pattern in the tissues (**Figure 4A** and **Figure 4B**). The control strain, PA14, showed the highest protease production (63.95 to 76.22 µg/mL in ASM), while clinical isolates SED 20 and SED 41 showed increased total proteases from 38.87 to 50.86 and 27.24 to 43.15 µg/mL, respectively, with meropenem treatment (**Figure 4A**). Interestingly, pyocyanin production varied between clinical strains with meropenem treatment. SED 20 and SED 43 infected tissues demonstrated decreased production by 68% in comparison to untreated, while SED 41 showed increased pyocyanin secretion by 162% with exposure to meropenem (**Figure 4B and Table S2)**. Surprisingly, ciprofloxacin treatment also showed a significantly higher pyocyanin by 116% (**Figure 4B and Table S2**). Proteases and pyocyanin concentrations in tissue and surrounding ASM, and total fold increases associated with antibiotic treatments are summarised in **Table S1** and **Table S2**, respectively.

**Figure 4.**
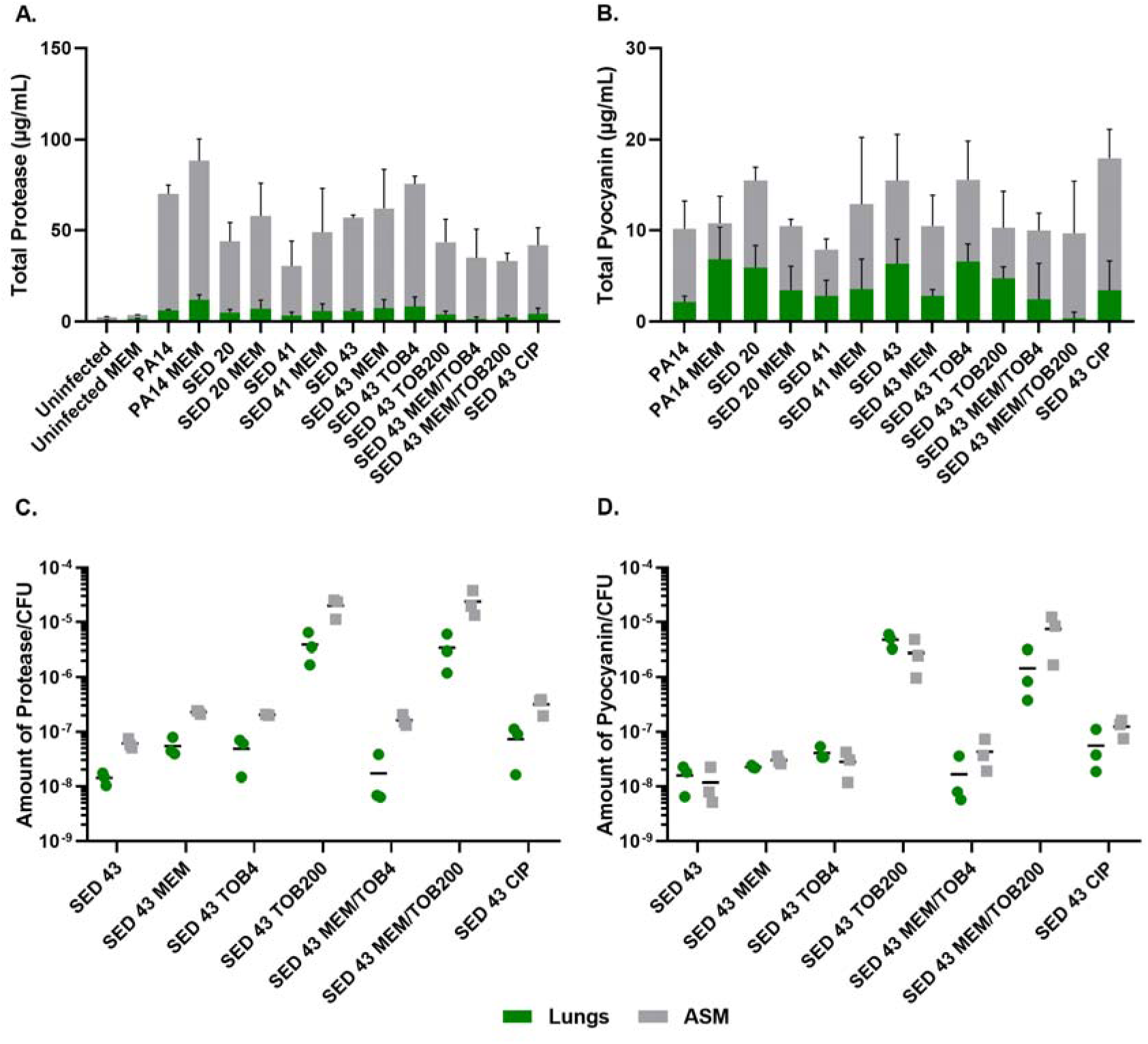
Using the EVPL model for understanding bacterial virulence with and without AB in comparison to control lab strains. **A.** Total protease, **B.** Total pyocyanin, **C.** The amount of protease/CFU, **D.** The amount of pyocyanin/CFU.

ANOVA showed a significant effect of strain (F_4,20_ = 6.798, p < 0.01) and meropenem treatment (F_1,20_ = 6.519, p < 0.05) on proteases production in the tissues, and the effect of meropenem did not differ between strains (strain*treatment interaction F_4,20_ = 0.825, p = 0.52). In the surrounding ASM, similar effects were observed with strain (F_4,20_ = 20.92, p < 0.001) and meropenem treatment (F_1,20_ = 3.193, p < 0.1). Total pyocyanin produced in the tissues and ASM was also significantly different between strains (F_4,20_ = 5.03, p < 0.01 and F_4,20_ = 6.40, p < 0.01, respectively), and the effect of meropenem significantly differed between strains (strain*treatment interaction F_4,20_ = 3.31, p < 0.05).

To further assess the potential effects on virulence of different clinically-relevant antibiotics, we exposed EVPL biofilms of strain SED 43 to ciprofloxacin, tobramycin and a combination of meropenem and tobramycin, in ASM. Interestingly, SED 43 showed a greater total production of proteases, in ASM, compared with the other clinical isolates (50.91 µg/mL), and slightly increased total proteases with meropenem (54.85 µg/mL, 109%) and tobramycin at low concentration (67.24 µg/mL, 133%). Both treatments led to comparable decreases in bacterial load (**Figure S5**). The increased bacterial death with ciprofloxacin and tobramycin, at high concentration, (**Figure S5**) correlated with decreased total proteases of 37.55 µg/mL (74%) and 39.7 µg/mL (77%), respectively. Combination treatment of tobramycin at 4 or 200 µg/mL with meropenem (64 µg/mL) showed a total protease of 33.83 (62%) and 31.07 µg/mL (59%) (**Figure 4A and Table S2**), respectively.

Because the total concentrations of proteases and pyocyanin discussed above are a function of both altered cellular production levels and altered cell numbers, we then normalised total proteases and pyocyanin concentrations by bacterial counts, to determine how antibiotic exposure affected per-cell production. The amount of proteases/cfu and pyocyanin/cfu measured in the tissues were slightly lower or equal to that of the surrounding ASM, respectively (**Figure 4C** and **4D**). Exposure to meropenem (64 µg/mL), ciprofloxacin (64 µg/mL) and tobramycin (4 µg/mL) slightly increased the production of proteases and pyocyanin by bacterial cfu, while a significant increase was found with tobramycin (200 µg/mL) and a combination of meropenem/tobramycin (64/200 µg/mL) (**Figure 4C** and **4D**). The latter treatments have a greater effect on bacterial load (**Figure S5**). The total amount of tissue and ASM proteases/cfu and pyocyanin/cfu is shown in **Figure S6**.

ANOVA analysis showed a significant effect of antibiotic treatment on the amount of protease/cfu (tissue F_6,13_ = 11.27, p < 0.001) and (surrounding ASM F_6,13_ = 37.09, p < 0.001), and production of pyocyanin (tissue F_6,13_ = 25.61, p < 0.001) and (surrounding ASM F_6,13_ = 17.88, p < 0.001).

## Discussion

In CF, chronic *P. aeruginosa* infections are characterised by the mucoid phenotype, which can adversely affect the individuals’ pulmonary function increasing mortality rates. Therefore, oral and nebulised ciprofloxacin and colistin, respectively, are administered at early infection stages to reduce the risk of chronic infection and during chronic infections to eradicate *P. aeruginosa* [18]. Un-cleared infections and moderate to severe exacerbation cases are treated with intravenous anti-pseudomonal antibiotics such as ceftazidime, meropenem and tobramycin in combination with β-lactams [18]. Pharmacokinetic characterisation of oral ciprofloxacin administration in CF has shown a C_max_ of 2.3 µg/mL in sputum [19]. The administration of a single intravenous dose of meropenem (1 gm) has been shown to achieve a bronchial secretion concentration of 0.53 µg/mL [20]. Intravenous tobramycin has been shown to lead to a sputum concentration of 68 µg/mL and to show a bactericidal effect at a sputum concentrations of 25x-MIC [21].

In this study, we compared the antibiotic susceptibility of selected *P. aeruginosa* strains using current diagnostic methods and our previously developed EVPL biofilm model. The increase in bacterial resistance profile with the 1 day biofilm Calgary device in comparison to standard MIC had been previously reported [7]. However, the Calgary device still does not represent *in vivo* biofilms. Exposure to ciprofloxacin and meropenem at concentrations higher than MBEC values and reported sputum concentrations led to a decrease of bacterial load by only 1-2 logs (**Figure 2**), which may be attributed to the formation of denser or mature biofilm in the EVPL model [17]. The failure to eradicate *P. aeruginosa* biofilms in the EVPL model, with such high concentrations, may be closer to the *in vivo* effect of these antibiotic treatments, demonstrating the need to employ CF representative diagnostics to better reflect on the antibiotics’ inhibitory effect.

Antibiotic tolerance was also affected by the use of different media. Besides the difference in planktonic and biofilm based models, Kirchner *et al.* demonstrated increased biofilm inhibitory concentrations, for most tested *P. aeruginosa* strains, when assessed in ASM compared with planktonic-based MIC in LB medium [22]. Davies *et al.* also showed the increase in bacterial heterogeneity, population diversity and antibiotic resistance of *P. aeruginosa* in ASM [23]. Therefore, we believe it is a more accurate model to show the effect of antibiotic treatment in CF is by testing for antibiotic susceptibility in ASM rather than general laboratory medium. This was alarmingly poorer than in MHB (**Figure 3**).

The work also indicated the potential *in vivo* effect of exposure to different antibiotics, in ASM, onto the virulence of *P. aeruginosa* to understand the clinical implications of chosen antibiotics. We focused on two main virulence factors of *P. aeruginosa*: proteases and pyocyanin production. Production of proteases is triggered by the quorum sensing system to degrade vital host proteins and antibodies. In CF lungs, proteases have been shown to cause a severe inflammatory response leading to pulmonary damage [24], and were detected in sputum during exacerbation [25]. Pyocyanin is also regulated by the quorum sensing system. It is a redox molecule that generates reactive oxygen species to induce oxidative stress in host cells, leading to cell damage and lysis. *P. aeruginosa* is protected from these reactive oxygen species by its own catalases [24]. Previous studies had estimated the concentration of pyocyanin in sputum of as high as 16.5 µg/mL [26], similar to the detected values in **Figure 4B**. The antibiotic recovery plates of *P. aeruginosa* following Calgary biofilm susceptibility testing did not show any production of pyochelin, pyoverdine or pyocyanin (**Figure S2**), but following exposure to antibiotics in the EVPL model, bacterial expression of proteases and pyocyanin was increased (**Figure 4**). This highlights the aggressiveness of *P. aeruginosa* infection in CF.

## Conclusion

Bacteria causing biofilm infections are often assessed for their antibiotic resistance profile using standard planktonic MIC methods or simple biofilm platforms such as the Calgary device. These do not represent the environment bacteria inhabit *in vivo*, giving misleading results. The current gap in clinical outcomes and standard susceptibility testing results is a very clear evidence. Our results, taken from an *ex vivo* animal tissue model using host-mimicking growth medium are consistent with increased antibiotic tolerance in *in vivo* biofilms. It is possible that current antibiotic prescribing could not only fail to eradicate biofilm load, but also worsen lung conditions by increasing expression of virulence factors by surviving bacteria, which requires immediate action to help eradicate biofilm infections. Thus, further work with clinical samples will be required to determine the effect of antibiotic treatment on bacterial load, lung function and possibly exacerbations.

## Acknowledgment

We thank Prof. Sophie Darch and Prof. Steve Diggle for CF isolates of *P. aeruginosa* and Prof. Leo Eberl for *P. aeruginosa* PA14. We would also like to acknowledge Cerith Harries and Caroline Stewart for the use of the media preparation facilities within the School of Life Sciences, Warwick University. This work was funded by a Medical Research Council New Investigator Research Grant to FH (MR/R001898/1).

## Author contributions

MMH and FH contributed to the concept of the study. MMH designed, performed and analysed experiments as well as wrote the manuscript. NH helped in designing and performing some of the virulence assay experiments. ES performed pilot work. All authors revised and approved the manuscript.

