## Supplementary material for "Predicting antibiotic-associated virulence of *Pseudomonas aeruginosa* using an *ex-vivo* lung biofilm model": Figure S1, S2, S3, S4, S5, S6, Table S1, Table S2

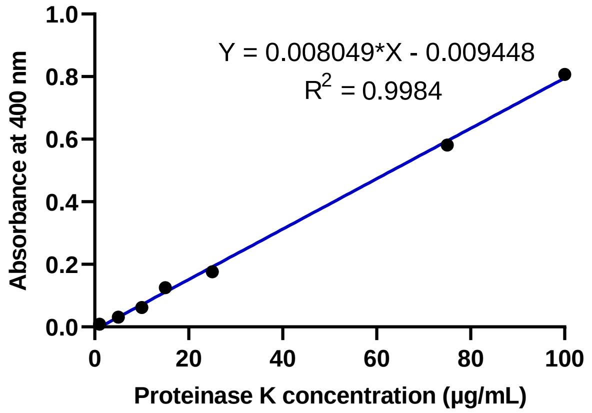


**Figure S1. Proteinase K standard curve.** Proteinase K standard curve was used to calculate the concentration of total protease produced by *P. aeruginosa* using the azocasein assay.

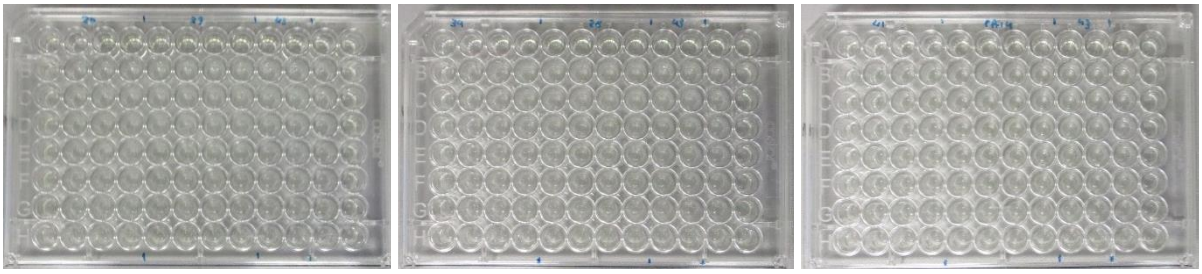

**Figure S2. MBEC recovery plates at 6 hour incubation of all tested bacterial strains.** There was no visual or plate reader detection of pyochelin, pyoverdine or pyocyanin.

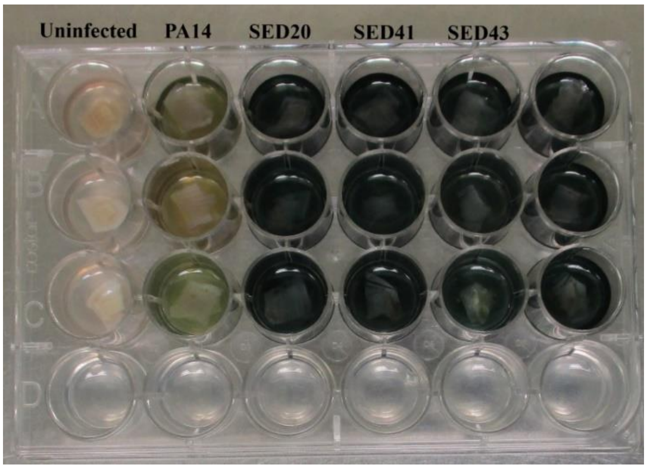


**Figure S3.** Uninfected and *P. aeruginosa* infected EVPL tissues after 7 days of biofilm formation.

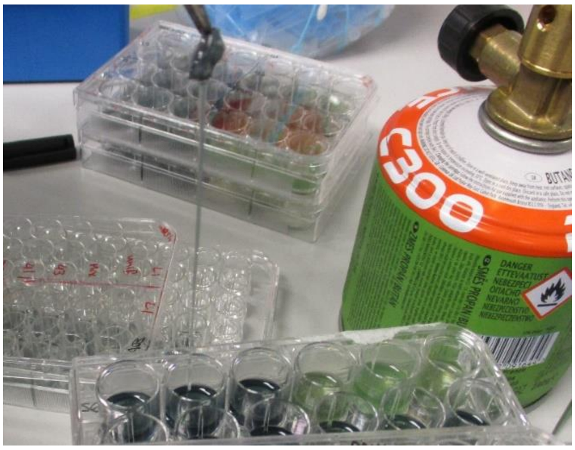


**Figure S4.** Mucoid phenotype of clinical strains after biofilm formation for 7 days.

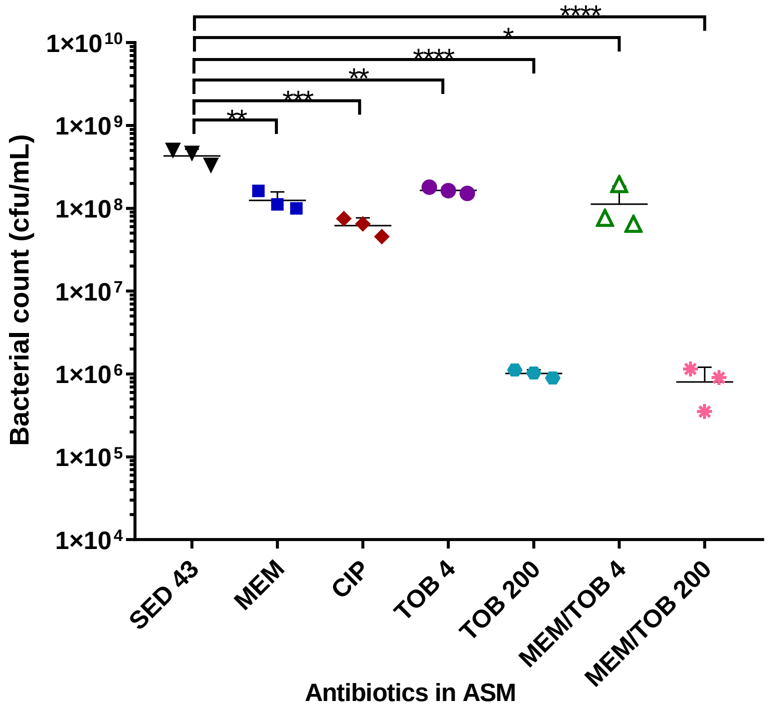


**Figure S5.** Bacterial load in the EVPL with different antibiotics in ASM.

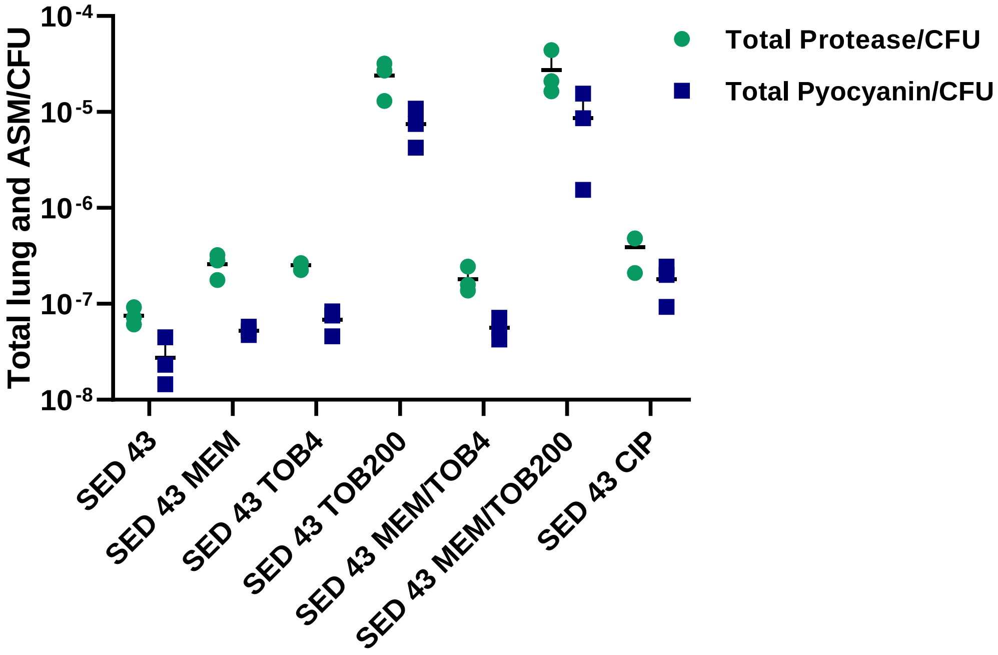


**Figure S6.** Total amount of produced protease and pyocyanin/cfu in both tissue and ASM.

**Table S1.** The concentration of protease and pyocyanin produced in the tissues and surrounding ASM.

| Strains/conditions | Concentration of (µg/mL) | | | |
|---|---|---|---|---|
| | Protease | | Pyocyanin | |
| | Tissue | ASM | Tissue | ASM |
| **Uninfected** | 0.54±0.34 | 1.91±0.21 | NA | NA |
| **Uninfected MEM** | 1.49±0.19 | 2.01±0.2 | NA | NA |
| **PA14** | 6.00±0.62 | 63.95±4.95 | 2.11±0.7 | 8.05±3.07 |
| **PA14 MEM** | 12.12±2.73 | 76.22±12.02 | 6.85±3.48 | 3.94±2.96 |
| **SED 20** | 4.97±1.63 | 38.88±10.52 | 5.93±2.42 | 9.56±1.44 |
| **SED 20 MEM** | 7.04±4.80 | 50.86±18 | 3.45±2.62 | 7.02±0.75 |
| **SED 41** | 3.27±1.92 | 27.24±13.61 | 2.82±1.72 | 5.11±1.10 |
| **SED 41 MEM** | 5.76±3.94 | 43.15±24.15 | 3.57±3.32 | 9.28±7.38 |
| **SED 43** | 5.87±0.77 | 50.91±1.66 | 6.33±2.69 | 9.15±5.08 |
| **SED 43 MEM** | 7.22±4.89 | 54.85±21.47 | 2.81±0.71 | 7.67±3.38 |
| **SED 43 TOB 4** | 8.24±5.37 | 67.24±4.40 | 6.65±1.85 | 8.92±4.27 |
| **SED 43 TOB 200** | 3.75±1.97 | 39.70±12.58 | 4.76±1.25 | 5.51±4.02 |
| **SED 43 MEM/TOB 4** | 1.42±0.98 | 33.83±15.35 | 2.65±3.77 | 7.50±1.94 |
| **SED 43 MEM/TOB 200** | 2.20±1.17 | 31.07±4.40 | 0.77±0.34 | 9.29±5.74 |
| **SED 43 CIP** | 4.22±3.04 | 37.55±9.61 | 3.43±3.27 | 14.52±3.18 |

**Table S2.** Fold increase in total concentration of protease and pyocyanin produced under different treatments. Blue and orange coloured values are indicating an increased and decreased percentage, respectively, in comparison to non-antibiotic treated tissues.

| Conditions | Fold increase | |
|---|---|---|
| | Total protease | Total pyocyanin |
| PA14 | 100 | 100 |
| PA14 MEM | 126 | 106 |
| SED 20 | 100 | 100 |
| SED 20 MEM | 132 | 68 |
| SED 41 | 100 | 100 |
| SED 41 MEM | 160 | 162 |
| SED 43 | 100 | 100 |
| SED 43 MEM | 109 | 68 |
| SED 43 TOB4 | 133 | 101 |
| SED 43 TOB200 | 77 | 66 |
| SED 43 MEM/TOB4 | 62 | 66 |
| SED 43 MEM/TOB200 | 59 | 65 |
| SED 43 CIP | 74 | 116 |
